# Human single-neuron recordings reveal population coding of attentional dynamics during naturalistic movie viewing

**DOI:** 10.64898/2026.09.08.748346

**Authors:** Carlo Cerquetella, Salman E. Qasim

**Affiliations:** Rutgers University, Robert Wood Johnson Medical School, Neurosurgery Department, New Brunswick, United States

**Keywords:** eye tracking, single-neuron recordings, naturalistic viewing, canonical correlation, intracranial electrophysiology, distributed population activity

## Abstract

Our eyes move constantly. Where they land is shaped by internal cognitive state, making eye movements a rare, non-invasive window onto that state. However, how moment-to-moment eye-movements map onto human neural population activity during naturalistic viewing remains poorly characterized. We analyzed a public intracranial dataset in which 14 neurosurgical patients watched an eight-minute movie during simultaneous eye-tracking and neuronal activity recording. Single-neuron activity from 814 neurons across anterior cingulate cortex, pre-supplementary motor area, amygdala, hippocampus, and ventromedial prefrontal cortex was pooled into a single distributed population and related to a joint eye-movement state comprising pupil size, saccade rate and fixation duration. Using a cross-validated, multivariate analysis of population activity we found group-level coupling between the neural population and the joint eye-movement state (mean held-out canonical r = 0.145; one-sample t-test p = 0.0003; 9 of 14 subjects individually significant). The coupling was carried principally by saccade dynamics (r = 0.141) and pupil size (r = 0.094), with fixation duration contributing only marginally (r = 0.041). At the single-neuron level the picture differed: more neurons were eye-coupled than expected by chance, yet so weakly that almost none survived correction for multiple comparisons. Read out jointly, the same weakly coupled cells cohered into one reliable population dimension, and no single region’s removal significantly reduced it. This pattern is more consistent with a distributed, redundant organization than with a small set of strongly coupled cells. In summary, eye-tracking offers a non-invasive window onto population-level neural states in humans.

## 1. Introduction

During natural vision the eyes are never still: humans redirect gaze three to four times per second, and where and how they look is shaped moment to moment by the internal cognitive and attentional state (Yarbus, 1967; Hayhoe and Ballard, 2005). Eye movements are thus a rich, continuous and non-invasive readout of that state. The three canonical variables of eye movements indicate partly dissociable processes: pupil size tracks arousal and the activity of ascending neuromodulatory systems (Aston-Jones and Cohen, 2005; Reimer et al., 2014; McGinley et al., 2015; Joshi et al., 2016); saccades reflect the active, attentional sampling of the scene (Corbetta and Shulman, 2002; Krauzlis et al., 2013); and fixation dynamics reflect the stable engagement and micro-behavior of the fovea on the currently attended target (Rucci and Poletti, 2015). These measures overlap without coinciding: they covary because a common behavioral state drives them, yet each retains dynamics the others do not capture. If that partial dissociation is mirrored in the brain, different aspects of the eye-movement state should couple to neural activity to different degrees, and potentially through partly different populations. Which of them the brain tracks, and where, is therefore an empirical question and cannot be settled by choosing a proxy in advance.

Naturalistic movie viewing engages these dynamics continuously and under ecological conditions, making it a powerful paradigm for studying how brain activity relates to the eye-movement state (Berg et al., 2009; Hasson et al., 2004). Most existing links between eye movements and neural activity in humans, however, come from scalp electroencephalography, functional MRI, or from animal models; direct evidence at the level of individual human neurons, recorded during free viewing of a naturalistic stimulus, remains rare. Human single-neuron recordings from intracranial depth electrodes, implanted for clinical seizure localization in patients with drug-resistant epilepsy, provide a unique opportunity to close this gap: they combine the activity of individual neurons, at a temporal resolution not limited by hemodynamics, with simultaneous behavioral monitoring (Rutishauser et al., 2006; Mormann et al., 2008). These electrodes routinely sample medial-temporal and medial-frontal structures—the amygdala (Amy), hippocampus (Hip), anterior cingulate cortex (ACC), pre-supplementary motor area (preSMA) and ventromedial prefrontal cortex (vmPFC)—that are implicated in arousal, salience, evaluation, oculomotor control and memory-guided viewing (Corbetta and Shulman, 2002; Aston-Jones and Cohen, 2005; Rutishauser et al., 2006; Rushworth et al., 2011; Bartra et al., 2013; Krauzlis et al., 2013). Whether, and at what grain, the population activity of these regions tracks the ongoing eye-movement state during naturalistic vision has not been characterized at the level of individual neurons.

Testing for such coupling rigorously is non-trivial. First, both neural firing and eye-movement statistics carry slow, shared drift, from fluctuating arousal, adaptation, and non-stationarity in the recordings, that can generate spurious correlations (Yule, 1926; Granger and Newbold, 1974). Second, the three canonical eye-movement measures are correlated expressions of a common behavioral state, so testing any one of them in isolation privileges a proxy chosen a priori and leaves the structure they share untested. Finally, mass-univariate single-neuron statistics ask whether individual cells track a feature, but cannot reveal whether many weak, scattered per-neuron relationships cohere into a single, coordinated population-level relationship. Whether the brain’s tracking of the eye-movement state is a property of individual neurons or of the population is not a technical detail, but the substantive question, and answering it requires a multivariate, population approach such as canonical correlation analysis (CCA; Hotelling, 1936).

Here we ask two questions. First, whether the distributed single-neuron population and the joint eye-movement state are coupled during continuous naturalistic movie viewing. Second, and more consequentially, at what level of organization any such coupling exists: whether it is carried by individual neurons tuned to the eye-movement state, or whether it is a property of the population that appears only when many weakly related cells are read out together. Using a publicly available multimodal intracranial dataset (Keles et al., 2024), we related the pooled firing of neurons recorded across five medial-frontal and limbic regions to a joint eye-movement state (pupil size, saccade rate, fixation duration) with a leak-free, drift-controlled, cross-validated CCA and a circular-shift surrogate null. We then examined the coupling with region loadings, leave-one-region-out analyses and single-neuron univariate tests, and we controlled for the number of recorded neurons. We find a reliable coupling that is carried principally by saccades and pupil size and that is distributed and redundant across the population, with no single region strictly necessary: a coordinated, population-level neural activity correlated to the eye-movement state that is invisible to single-neuron statistics alone.

## 2. Materials and methods

### 2.1 Participants and dataset

We analyzed the publicly available multimodal dataset DANDI:000623 (Keles et al., 2024; CC-BY-4.0), in which 16 patients with drug-resistant epilepsy underwent intracranial depth-electrode recording (single units and local field potentials) while watching an eight-minute short movie (640 × 480 pixels, 25 frames per second) with simultaneous eye-tracking. All original data were collected under the ethical approvals and informed-consent procedures reported in the source publication; the present study is a secondary analysis of de-identified data and involved no new data collection. Two subjects were excluded a priori for insufficient neural yield: sub-CS41 (7 units) and sub-CS49 (12 units), both below the inclusion threshold of ≥ 15 units across all recorded regions. The threshold was set on unit count alone, independently of any coupling outcome. The final cohort comprised 14 subjects (sub-CS42, sub-CS43, sub-CS44, sub-CS47, sub-CS48, sub-CS51, sub-CS53, sub-CS54, sub-CS55, sub-CS56, sub-CS57, sub-CS58, sub-CS60, sub-CS62).

Single-unit spike times and eye-tracking are referenced to movie onset (t = 0) and were analyzed within the movie-encoding window [0, 478.85 s]. Spike counts were pooled across all recorded units, spanning five bilateral target regions: ACC, Amy, Hip, preSMA and vmPFC, and smoothed into 25 Hz firing-rate traces with a Gaussian kernel (σ = 250 ms). Recording coverage across the cohort is summarized in Fig. 1A. All signals were placed on a common 25 Hz grid (one bin = one movie frame = 40 ms, Fig. 1B).

**Fig. 1.**
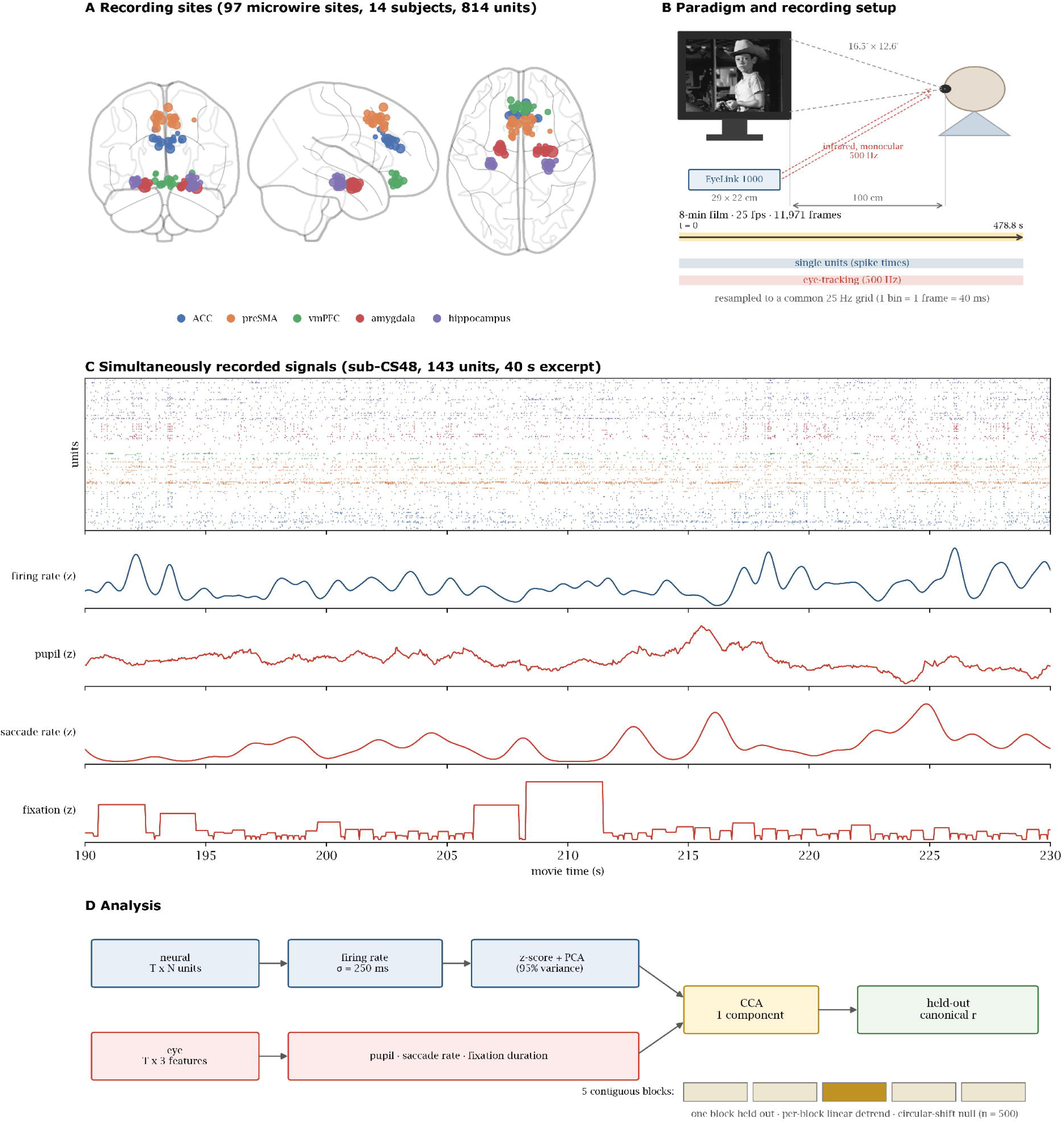
Schematic overview. (A) Recording sites. All 97 distinct microwire locations contributing units to the analyzed cohort (14 subjects, 814 units: ACC 132, preSMA 177, vmPFC 89, amygdala 260, hippocampus 156 units), plotted in MNI space and colored by target region. (B) Paradigm and recording setup. Subjects watched an eight-minute film (25 frames per second) presented at 29 × 22 cm from a viewing distance of 100 cm (16.5° × 12.6° of visual angle), while monocular gaze position and pupil size were recorded at 500 Hz with an EyeLink 1000 and single-neuron activity was recorded from intracranial depth electrodes. The encoding window [0, 478.85 s] was analyzed, and both variables were placed on a common 25 Hz grid (one bin = one movie frame = 40 ms). The image on the monitor is a 1961 NBC publicity for the episode used as the stimulus (“Bang! You’re Dead”, Alfred Hitchcock Presents; public domain). (C) Simultaneously recorded signals, 40 s excerpt (190–230 s) from sub-CS48 (143 units). Top: spike raster, one row per unit, colored by region as in A. Bottom: pooled firing rate (Gaussian σ = 250 ms, z-scored) and the three eye-movement features (pupil size, saccade rate and fixation duration) on the shared movie-time axis. (D) Pooled spike counts were converted to firing rates, then standardized and reduced by PCA (95% variance) within each training fold, and related to the joint eye-movement state by a one-component CCA. Cross-validation used five contiguous blocks with a per-block linear detrend, and significance was assessed against a circular-shift surrogate null (n = 500 shifts), yielding one held-out canonical correlation per subject.

### 2.2 Eye-tracking acquisition and eye-movement features

Eye-tracking was acquired by the original investigators; full details are given in Keles et al. (2024), and we summarize here those relevant to the present analysis. Monocular gaze position and pupil size were recorded at 500 Hz with an EyeLink 1000 (SR Research Inc.) using infrared corneal reflection, with a sticker used to track head position. The movie was presented with the Psychophysics Toolbox in MATLAB at 25 frames per second, subtending 29 × 22 cm at a viewing distance of 100 cm (16.5° × 12.6° of visual angle). Pupil size was recorded as the number of pixels within the pupil contour and is therefore in arbitrary units; it was z-scored within subject, so only relative fluctuations enter the analysis. Fixations, saccades and blinks were classified with the EyeLink’s built-in event-detection algorithms and are distributed with the dataset as event tables of onset times and durations; we used these as provided rather than re-detecting events from raw gaze. In the source dataset, gaze samples were missing for 7% ± 8% of the movie-watching period (mean ± S.D. across participants and runs) and individual gaze patterns correlated with a reference visual-salience map at r = 0.75 ± 0.12, indicating good overall tracking quality (Keles et al., 2024). The recording setup is shown schematically in Fig. 1B, and an excerpt of the simultaneously recorded neural and eye-movement signals in Fig. 1C.

Three eye-movement features were constructed at 25 Hz and z-scored within subject. For pupil size, samples were marked missing if the pupil ≤ 0 (approximately 10% of samples), if they fell within an author-provided blink interval padded by ±50 ms, or if they remained > 3 SD from the median of the surviving samples; the resulting gaps were linearly interpolated at the native 500 Hz rate, and the cleaned trace was z-scored and resampled onto the 25 Hz grid. The saccade feature was a binary saccade-occupancy trace convolved with a Gaussian kernel (σ = 12 bins = 480 ms). The fixation feature assigned each fixation’s duration to every bin it occupied. Features were analyzed jointly (all three simultaneously) for the primary test, and individually for feature attribution. The three measures are not independent: saccade and pupil responses are coordinated by shared midbrain orienting circuitry and covary trial by trial (Wang and Munoz, 2015, 2021), fixation duration is by construction the interval between successive saccades (Otero-Millan et al., 2008), and all three track arousal (Reimer et al., 2014; McGinley et al., 2015). What they jointly express, the eye-movement state, is the quantity of interest, and CCA estimates it directly, learning the combination of eye features best aligned with population activity rather than fixing a proxy in advance (Wang et al., 2020). Each feature’s separate contribution is recovered in the attribution analysis (Section 3.2).

### 2.3 Canonical correlation pipeline

Coupling between the neural population and the eye-movement state was quantified with a cross-validated CCA, the standard approach for relating a set of neural variables to a heterogeneous set of behavioral measures (Smith et al., 2015; Wang et al., 2020). Spike counts were first converted to firing rates by Gaussian smoothing (σ = 250 ms) applied once to the full recording; within each cross-validation fold the neural data were then standardized and reduced by principal component analysis retaining 95% of variance, with both the scaler and the PCA fit on the training fold only. A one-component CCA, regularized by the preceding PCA reduction, was then fit between the pooled principal-component matrix and the T × 3 joint eye-feature matrix, and coupling was measured as the held-out canonical correlation. Because smoothing precedes the split, samples within ∼3σ (0.75 s) of a fold boundary are marginally shared across folds; this is negligible relative to the ∼96 s block length and applies identically to the surrogate null. Cross-validation used contiguous five-fold block partitioning rather than interleaved folds, because interleaving interpolates slow drift across fold boundaries and inflates the null.

Both neural firing rates and eye-movement statistics drift slowly over an eight-minute recording, through electrode drift, adaptation, and gradual changes in arousal. Shared drift is a well-established source of spurious correlation between otherwise unrelated time series (Yule, 1926; Granger and Newbold, 1974). The circular-shift null does not by itself neutralize this, since the surrogates inherit the same slow structure. Drift was therefore controlled explicitly (primary analysis) by removing a per-block linear trend within each fold before CCA: a straight-line fit removes slow drift without the edge artifacts of Gaussian high-pass filtering and without smearing structure across the global fold boundary (Supplementary Fig. S1). Each block, including the held-out block, is detrended using only its own samples; this step is unsupervised and is applied identically to the observed data and to every surrogate. Together these steps define the temporal band in which coupling is estimated: the σ = 250 ms firing-rate kernel attenuates neural fluctuations above ≈ 0.5 Hz (half-power), the σ = 480 ms saccade-rate kernel above ≈ 0.3 Hz, and the per-block linear detrend removes fluctuations slower than roughly one cycle per fold (≈ 0.01 Hz). The null distribution was generated by circularly shifting the eye-feature block (n = 500 shifts), which preserves each feature’s autocorrelation and the cross-feature covariance while destroying the neural–behavioral alignment. Significance was reported as an empirical p (floor 1/501 ≈ 0.002). For feature attribution, each feature was analyzed in isolation under identical settings; because the target is then one-dimensional, the fit reduces to ridge regression and coupling is the held-out correlation between predicted and observed feature (equivalent to a regularized single-feature canonical correlation). Fig. 1D summarizes the pipeline.

### 2.4 Structure coupling analyses

Three complementary analyses established where the coupling is expressed, and at what grain. Region loadings were obtained by projecting the first neural canonical variate back onto individual neurons grouped by region, for each individually significant subject, and computing each region’s share of the total loading magnitude (shares sum to one within subject). These loadings come from a single fit on all timepoints (not cross-validated) under the global Gaussian high-pass drift control, and are therefore descriptive: they indicate how the shared dimension is distributed across regions, not whether any region is necessary.

Necessity was assessed by leave-one-region-out: each region’s units were removed from the pooled population and the pipeline re-run, and the paired difference Δr = r_full − r_drop was tested across subjects with a right-tailed one-sample t-test. A subject contributed to a region’s test only if it had units in that region and retained at least 15 units after removal, so the number of subjects varies across regions.

At the single-neuron level, each neuron’s smoothed firing rate was correlated with each z-scored feature (per-block linear detrend) using Spearman rank correlation, which is robust to the skewed firing-rate distribution, against a circular-shift surrogate null (n = 10,000). Per-unit significance was two-sided on |r| at raw empirical p < 0.05, with Benjamini–Hochberg correction applied across all units within a feature. The excess over the 5% chance level was tested with a pooled binomial test per region and with a nesting-aware per-subject test; because the binomial pools units across subjects and so ignores within-subject dependence (anti-conservative), the per-subject test is the inferential statement.

### 2.5 Population-size control and statistics

To rule out a trivial sampling explanation (subjects with more neurons yielding higher canonical correlations) we regressed a per-subject canonical r² against unit count across the 14 subjects. Group inference used one-sample t-tests on the effect (observed r − null mean), reported two-tailed, alongside right-tailed Wilcoxon signed-rank tests; the leave-one-region-out and per-subject fraction tests were right-tailed. Between-region differences in per-unit |r| used cluster-robust ordinary least squares accounting for neuron-within-subject nesting, complemented by Kruskal–Wallis and Dunn tests. Multiple comparisons across features and regions were controlled with the Benjamini–Hochberg false discovery rate (FDR; Benjamini and Hochberg, 1995) at α = 0.05. Analyses were run with fixed random seeds.

## 3. Results

### 3.1 The distributed neural population is coupled to the joint eye-movement state

Across the 14-subject cohort, the pooled neural population and the joint eye-movement state shared a significant canonical dimension. The mean held-out canonical correlation was r = 0.145, significantly above the circular-shift null at the group level (one-sample t-test on the effect, observed r - null mean, p = 0.0003; Wilcoxon signed-rank p = 0.0006), and 9 of 14 subjects were individually significant (Table 1; Fig. 2). The eye-movement state is therefore reflected in a coordinated dimension of distributed single-neuron population activity during naturalistic viewing.

**Fig. 2.**
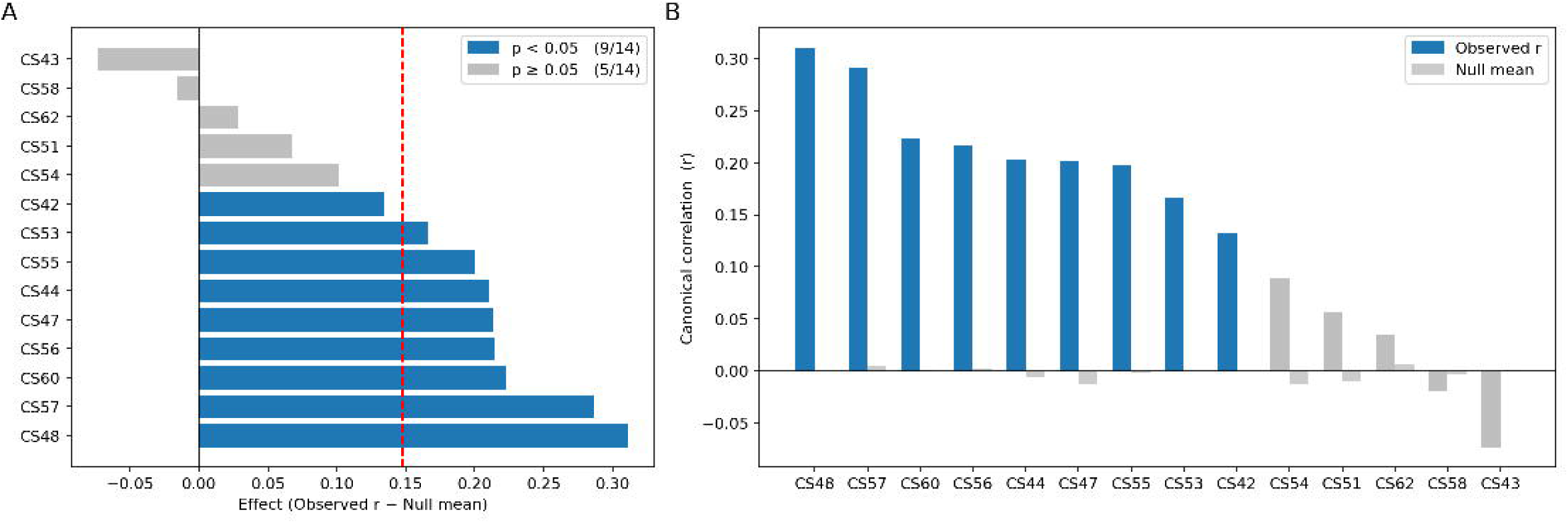
Per-subject neural–eye coupling effects (block CV + per-block linear detrend, n = 14). (A) Coupling effect (observed canonical r − null mean) per subject, sorted by effect size; blue = individually significant (circular-shift p < 0.05), grey = not significant; dashed red line = group mean effect. (B) Observed canonical r (colored) versus circular-shift null mean (light grey) per subject. Group one-sample t: p = 0.0003, Wilcoxon p = 0.0006; 9/14 subjects individually significant (solid blue significant, solid grey non-significant subjects).

**Table 1.**
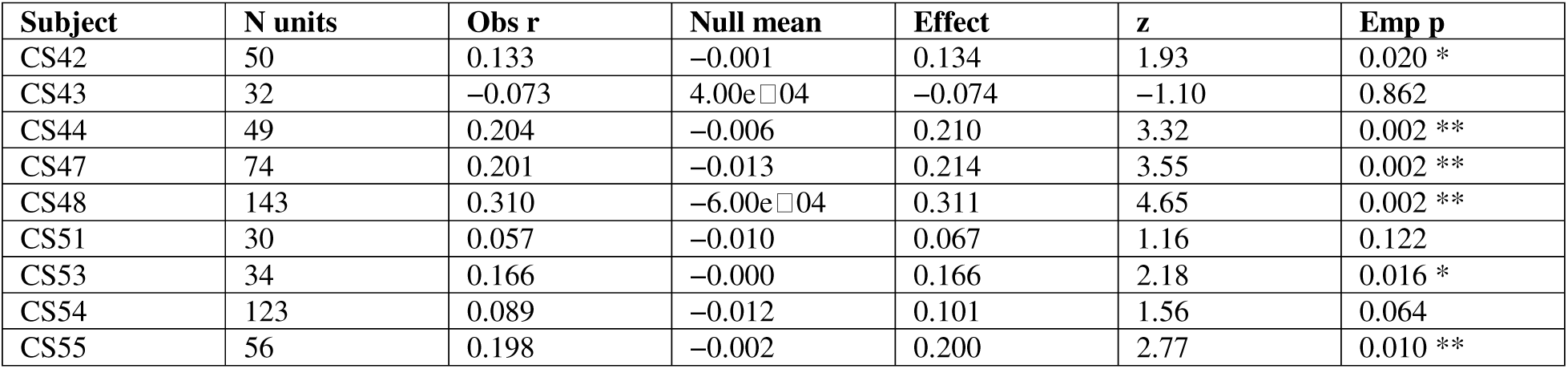
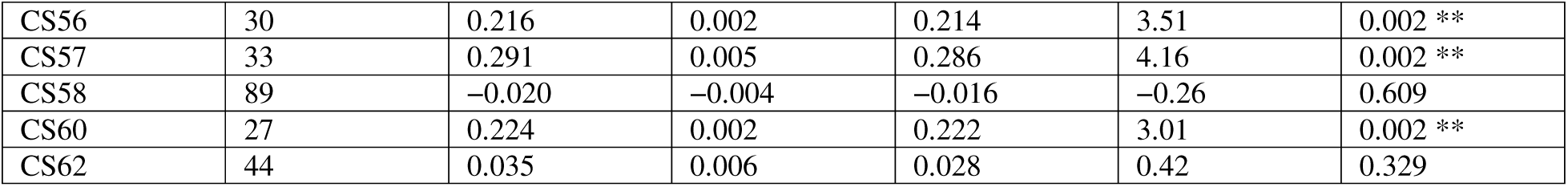
Per-subject primary coupling (block cross-validation + per-block linear detrend). Obs r = held-out canonical correlation; Effect = Obs r − null mean; p = empirical p against the circular-shift null (floor 1/501 ≈ 0.002). Mean null ≈ 0 (shifts are unbiased). 9 of 14 subjects are individually significant.

| Subject | N units | Obs $r$ | Null mean | Effect | $z$ | Emp $p$ |
| --- | --- | --- | --- | --- | --- | --- |
| CS42 | 50 | 0.133 | -0.001 | 0.134 | 1.93 | 0.020 * |
| CS43 | 32 | -0.073 | 4.00e-04 | -0.074 | -1.10 | 0.862 |
| CS44 | 49 | 0.204 | -0.006 | 0.210 | 3.32 | 0.002 ** |
| CS47 | 74 | 0.201 | -0.013 | 0.214 | 3.55 | 0.002 ** |
| CS48 | 143 | 0.310 | -6.00e-04 | 0.311 | 4.65 | 0.002 ** |
| CS51 | 30 | 0.057 | -0.010 | 0.067 | 1.16 | 0.122 |
| CS53 | 34 | 0.166 | -0.000 | 0.166 | 2.18 | 0.016 * |
| CS54 | 123 | 0.089 | -0.012 | 0.101 | 1.56 | 0.064 |
| CS55 | 56 | 0.198 | -0.002 | 0.200 | 2.77 | 0.010 ** |
| CS56 | 30 | 0.216 | 0.002 | 0.214 | 3.51 | 0.002 ** |
| CS57 | 33 | 0.291 | 0.005 | 0.286 | 4.16 | 0.002 ** |
| CS58 | 89 | -0.020 | -0.004 | -0.016 | -0.26 | 0.609 |
| CS60 | 27 | 0.224 | 0.002 | 0.222 | 3.01 | 0.002 ** |
| CS62 | 44 | 0.035 | 0.006 | 0.028 | 0.42 | 0.329 |

Two features of the canonical solution show that it reflects the joint eye-movement state rather than a single feature in disguise. First, the eye-side weights were distributed across features: at least two of the three carried more than 10% of the total weight in 12 of 14 subjects, and which feature dominated differed from subject to subject (Supplementary Table S1). Second, the way the features combined was consistent across subjects. The overall sign of a canonical pair is arbitrary, so we fixed it in each subject by making the saccade weight positive; pupil size then entered with the opposite sign in all 9 individually significant subjects, and in 12 of 14 subjects overall (binomial p = 0.004 and p = 0.013), whereas the fixation weight was negative in only 3 of the 9 and carried no consistent sign. The population dimension thus tracks the balance between saccadic sampling and pupil size, not their sum.

### 3.2 Saccades and pupil size carry most of the joint coupling

The joint test establishes that the neuronal population tracks the omnibus eye-movement state, but not how much each feature contributes. We therefore repeated the analysis with each feature in isolation (Table 2; Fig. 3). Saccade dynamics were the dominant carrier (mean r = 0.141, 8/14 subjects significant, group t p = 0.0003), followed by pupil size (r = 0.094, 7/14, p = 0.0005). Fixation duration was marginal on its own (r = 0.041, 3/14, p = 0.026) and drew on limbic rather than medial-frontal populations. Per-subject, per-feature values are given in Supplementary Table S2. Every saccade-significant subject was also significant jointly, and one subject (CS53) was individually significant on the joint state while showing essentially no saccade coupling (Supplementary Table S2). Together with the opposed saccade and pupil weights of Section 3.1, this indicates that the joint dimension is not recoverable by ranking the features one at a time.

**Fig. 3.**
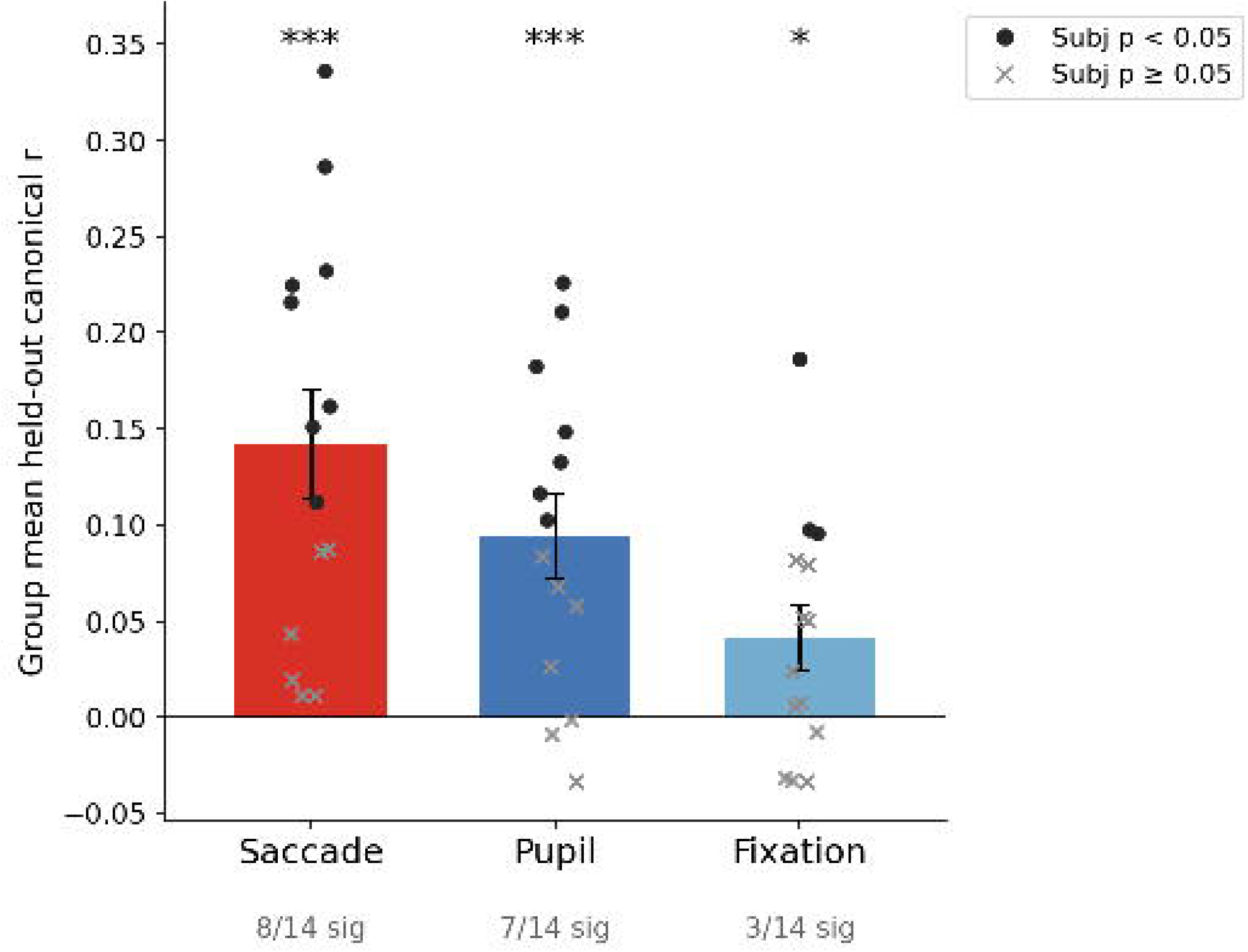
Per-feature attribution (individual-feature CCA, n = 14). Bars show group mean held-out canonical r ± SEM; individual subjects overlaid (filled circles = individually significant p < 0.05; crosses = p ≥ 0.05). Stars indicate group significance (*** p < 0.001; ** p < 0.01; * p < 0.05). Saccade is the dominant carrier (r = 0.141, 8/14), pupil is significant (r = 0.094, 7/14), and fixation is marginal (r = 0.041, 3/14).

**Table 2.**
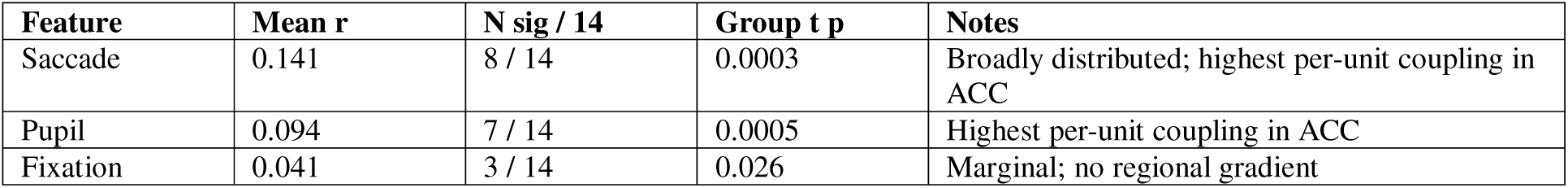
Per-feature attribution (single-feature canonical correlation, per-block linear detrend, block CV, 500-shift null). Group statistics are one-sample t-tests on the effect across 14 subjects.

| Feature | Mean r | N sig / 14 | Group t p | Notes |
| --- | --- | --- | --- | --- |
| Saccade | 0.141 | 8 / 14 | 0.0003 | Broadly distributed; highest per-unit coupling in ACC |
| Pupil | 0.094 | 7 / 14 | 0.0005 | Highest per-unit coupling in ACC |
| Fixation | 0.041 | 3 / 14 | 0.026 | Marginal; no regional gradient |

### 3.3 The coupling is distributed across regions and no single region is necessary

For each individually significant subject (n = 9), the first neural canonical variate was projected back onto neurons grouped by region (Table 3; Fig. 4). On average, preSMA and Amy each contributed about 31% of the coupling signal (together ≈ 62%), followed by ACC (≈ 20%), vmPFC (≈ 10%) and Hip (≈ 9%). The pattern was heterogeneous across subjects: Amy-dominant in CS47, preSMA-dominant in CS44; already hinting that no single region monopolizes the coupling.

**Fig. 4.**
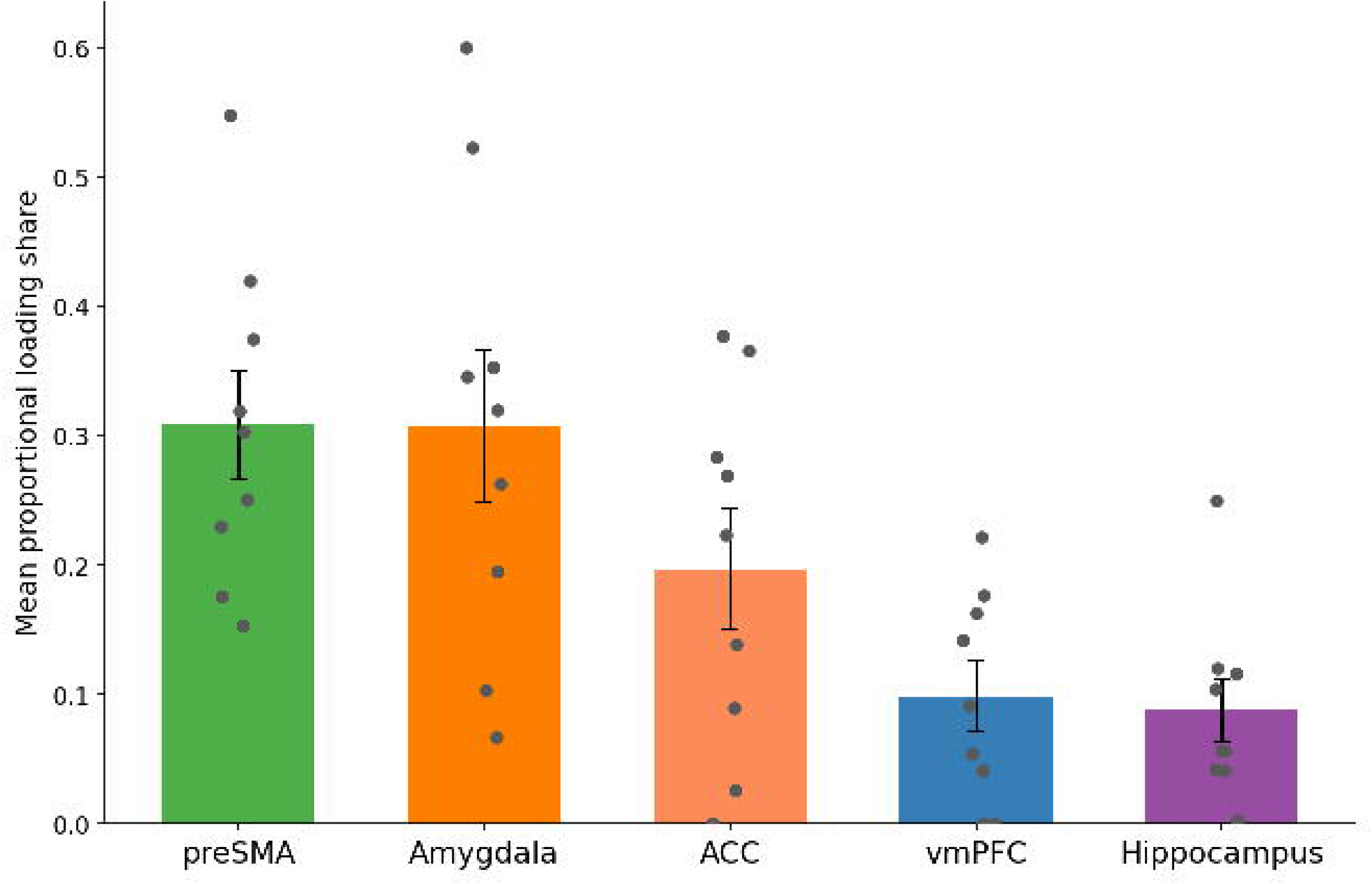
Proportional contribution of each region to the first neural canonical variate, averaged over the 9 individually significant subjects (mean ± SEM); dots show individual subjects. preSMA and Amy each contribute ≈ 31%; ACC ≈ 20%; vmPFC ≈ 10%; Hip ≈ 9%. The pattern is heterogeneous across subjects.

**Table 3.** Proportional region contribution to the first neural canonical variate (absolute loading share; rows sum to 1 within subject). Mean over the 9 individually significant subjects.

| Subject | ACC | Amy | Hip | preSMA | vmPFC | Top feature |
| --- | --- | --- | --- | --- | --- | --- |
| CS42 | 0.377 | 0.263 | 0.057 | 0.303 | 0.000 | Pupil |
| CS44 | 0.269 | 0.067 | 0.116 | 0.548 | 0.000 | Saccade |
| CS47 | 0.000 | 0.600 | 0.003 | 0.176 | 0.222 | Saccade |
| CS48 | 0.284 | 0.195 | 0.250 | 0.230 | 0.041 | Saccade |
| CS53 | 0.139 | 0.346 | 0.041 | 0.420 | 0.054 | Pupil |
| CS55 | 0.026 | 0.353 | 0.104 | 0.375 | 0.142 | Pupil |
| CS56 | 0.090 | 0.523 | 0.057 | 0.153 | 0.177 | Saccade |
| CS57 | 0.223 | 0.320 | 0.042 | 0.251 | 0.163 | Saccade |
| CS60 | 0.366 | 0.103 | 0.120 | 0.319 | 0.091 | Saccade |
| Mean (9 sig) | 0.20 | 0.31 | 0.09 | 0.31 | 0.10 | — |

To test necessity, each region’s units were removed from the pooled population and the pipeline re-run (Table 4; Fig. 5). No leave-one-region-out drop survived FDR correction (all q > 0.05). ACC and preSMA show the largest raw effects (Δr = +0.033, raw p = 0.020 and Δr = +0.027, raw p = 0.063 respectively) and indicate that these regions contribute to the coupling but are not required: the remaining population compensates. Exploratory per-feature leave-one-region-out (Supplementary Table S3) showed that saccade coupling leaned on ACC and preSMA, whereas fixation leaned on the Amy and Hip — complementary substrates not explained by saccade and pupil.

**Fig. 5.**
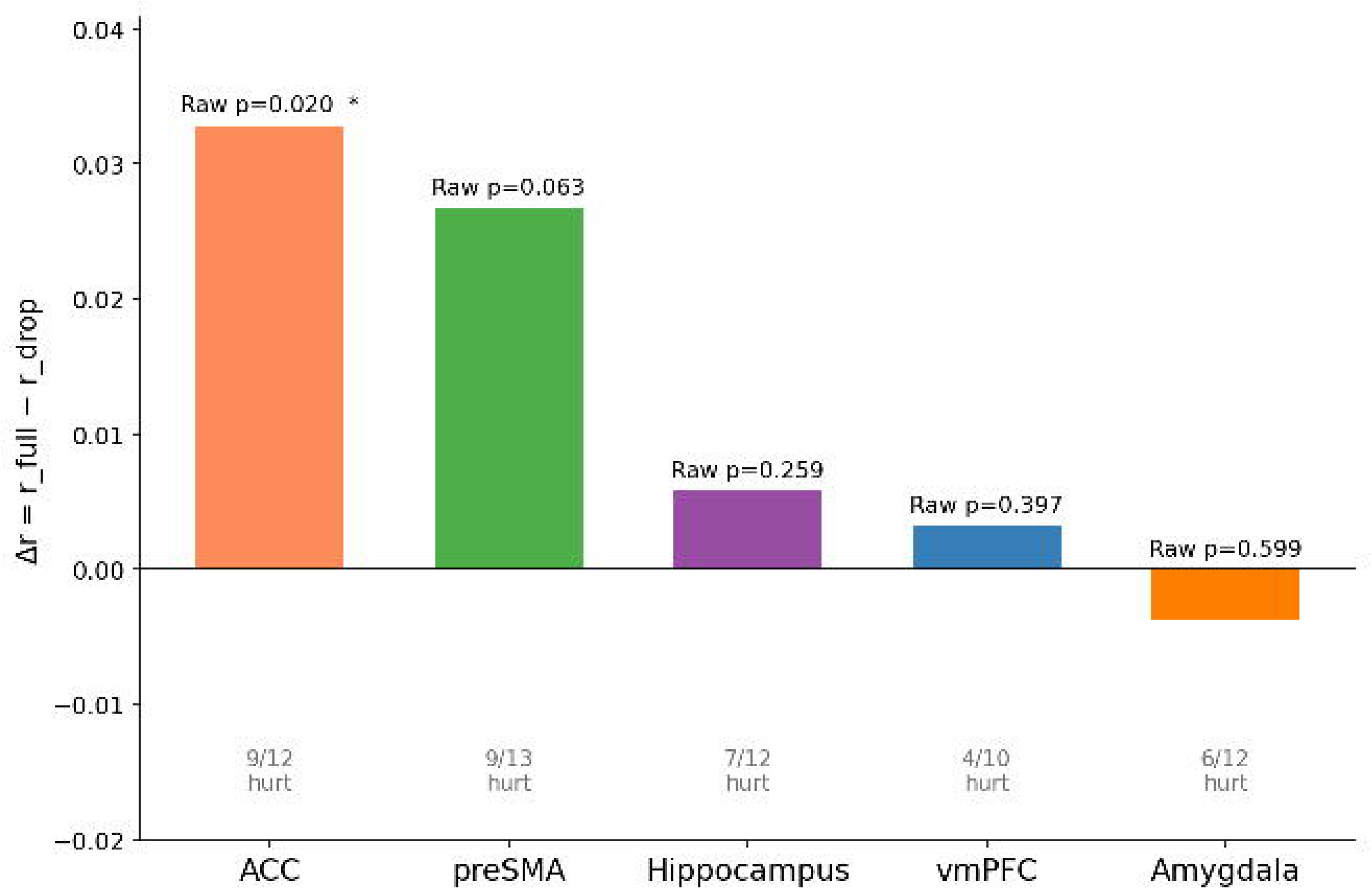
Leave-one-region-out. Bars show mean paired Δr = r_full − r_drop when each region’s units are removed; Δr > 0 means the region contributes when present. Annotated p-values are raw (uncorrected); * = raw p < 0.05. No region survives BH FDR (all q > 0.05), confirming a redundantly distributed organization with no single necessary region.

**Table 4.**
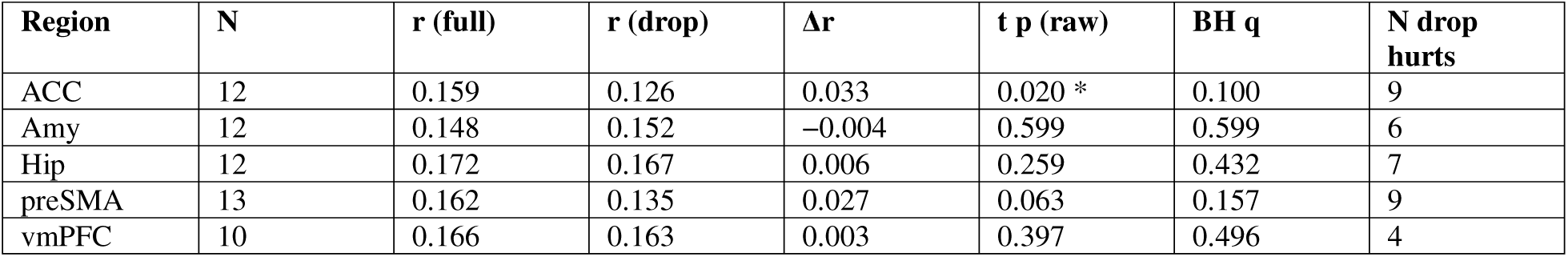
Leave-one-region-out (necessity). Δr = r_full − r_drop; one-sample t-test across subjects; BH FDR across 5 regions. No region survives correction. N = subjects contributing to that region’s test (units present in the region and ≥ 15 units remaining after its removal), which is why r (full) differs slightly across rows. “N drop hurts” = subjects for whom removing the region reduced coupling.

### 3.4 Single-neuron coupling is a distributed population excess

At the single-neuron level, coupling was frequent but individually weak (Table 5; Fig. 6). At the uncorrected threshold, saccade coupling was the most common and was elevated in every region, from 11.2% of units (vmPFC) to 18.9% (ACC) — about two to four times chance — with ACC highest, followed closely by Hip (16.7%) and preSMA (16.4%); pupil was weaker (highest in ACC, 15.2%), and fixation was only slightly above chance. After FDR correction, however, only a few neurons survived (saccade 28/814 units = 3.4%; pupil 6/814 = 0.7%; fixation 16/814 = 2.0%). Single-neuron coupling was therefore widespread but weak: a reliable excess of weakly coupled units, with few individual neurons surviving correction. This pattern is more consistent with a distributed, redundant organization than with a sparse set of strongly coupled driver neurons, and parallels the absence of any strictly necessary region (Section 3.3).

**Fig. 6.**
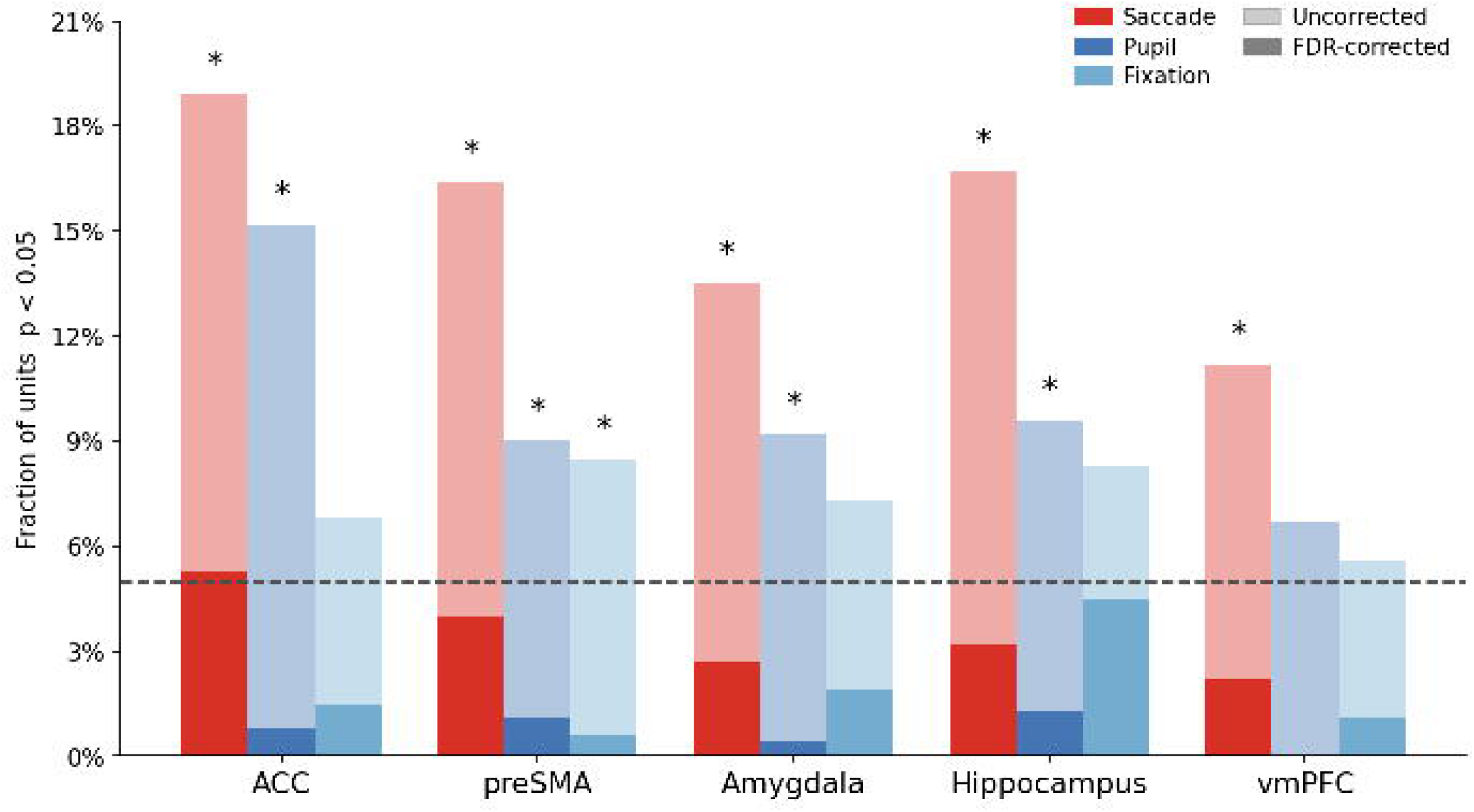
Single-neuron univariate coupling by region and feature. Pale bars = fraction of units coupled at the uncorrected threshold; solid bars = fraction surviving BH FDR correction; dashed line = 5% chance, which is the reference for the uncorrected bars only (under FDR control the expected null fraction among rejections is not 5%). Stars mark regions exceeding chance (pooled binomial, BH < 0.05). Raw saccade fractions sit roughly 2-4 x above chance while very few units survive correction: a distributed population excess of weakly coupled neurons, not a focal set of strongly coupled cells.

**Table 5.** Single-neuron univariate coupling by region and feature (Spearman rank correlation, circular-shift null). Frac sig = fraction of units at uncorrected empirical p < 0.05; chance ≈ 5%.

| Region | Feature | N units | Mean $\vert r\vert $ | Frac sig | N sig |
| --- | --- | --- | --- | --- | --- |
| ACC | fixation | 132 | 0.035 | 6.8% | 9 |
| ACC | pupil | 132 | 0.054 | 15.2% | 20 |
| ACC | saccade | 132 | 0.060 | 18.9% | 25 |
| Amy | fixation | 260 | 0.034 | 7.3% | 19 |
| Amy | pupil | 260 | 0.040 | 9.2% | 24 |
| Amy | saccade | 260 | 0.047 | 13.5% | 35 |
| Hip | fixation | 156 | 0.037 | 8.3% | 13 |
| Hip | pupil | 156 | 0.040 | 9.6% | 15 |
| Hip | saccade | 156 | 0.047 | 16.7% | 26 |
| preSMA | fixation | 177 | 0.033 | 8.5% | 15 |
| preSMA | pupil | 177 | 0.047 | 9.0% | 16 |
| preSMA | saccade | 177 | 0.049 | 16.4% | 29 |
| vmPFC | fixation | 89 | 0.037 | 5.6% | 5 |
| vmPFC | pupil | 89 | 0.043 | 6.7% | 6 |
| vmPFC | saccade | 89 | 0.047 | 11.2% | 10 |

Two population tests confirmed the excess is reliable. A pooled binomial test against the 5% chance level showed saccade coupling above chance in all five regions (ACC, preSMA, Hip and Amy q ≈ 0; vmPFC q = 0.025), pupil coupling above chance in ACC, Amy, Hip and preSMA (q ≤ 0.028) but not vmPFC, and fixation coupling above chance only in preSMA (q = 0.049). A nesting-aware per-subject test, comparing each subject’s coupled fraction with 5% (right-tailed), gave the same ordering: saccade mean 14.6% in 13 of 14 subjects (t p = 0.0001), pupil 9.5% in 12 of 14 (p = 0.0009), and fixation 7.7% in 11 of 14 (p = 0.022).

Between-region differences in per-unit |r| (cluster-robust OLS with neuron-within-subject nesting, BH-corrected; Table 6) were significant only for pupil (omnibus p = 0.029, uncorrected across the three features: ACC > amygdala and ACC > hippocampus, with Dunn and cluster-OLS in agreement). The saccade omnibus was not significant under nesting correction (p = 0.116); its apparent ACC lead appears only on the pooled Dunn test; fixation showed no regional differences (p = 0.80). Saccade coupling is thus broadly distributed rather than focal.

**Table 6.** Single-neuron between-region test. KW = Kruskal–Wallis; EMM = estimated marginal mean |r| from cluster-robust OLS (neuron-within-subject nesting; BH at the pairwise level). Only pupil shows a significant gradient (ACC > Amy, ACC > Hip).

| Feature | KW H | KW p | OLS omnibus p | EMM ACC | EMM Amy | EMM Hip | EMM preSMA | EMM vmPFC |
| --- | --- | --- | --- | --- | --- | --- | --- | --- |
| pupil | 15.08 | 0.005 | 0.029 * | 0.054 | 0.040 | 0.040 | 0.047 | 0.043 |
| saccade | 11.10 | 0.026 | 0.116 | 0.060 | 0.047 | 0.047 | 0.049 | 0.047 |
| fixation | 4.71 | 0.319 | 0.798 | 0.035 | 0.034 | 0.037 | 0.033 | 0.037 |

### 3.5 The coupling does not scale with the number of recorded neurons

Finally, we asked whether subjects with more neurons obtained higher canonical correlations by a law-of-large-numbers effect. Per-subject canonical r² showed no significant relationship with unit count (Pearson r = 0.196, p = 0.50, two-tailed; Fig. 7, Supplementary Table S4). The main result is therefore not explained by sample size.

**Fig. 7.**
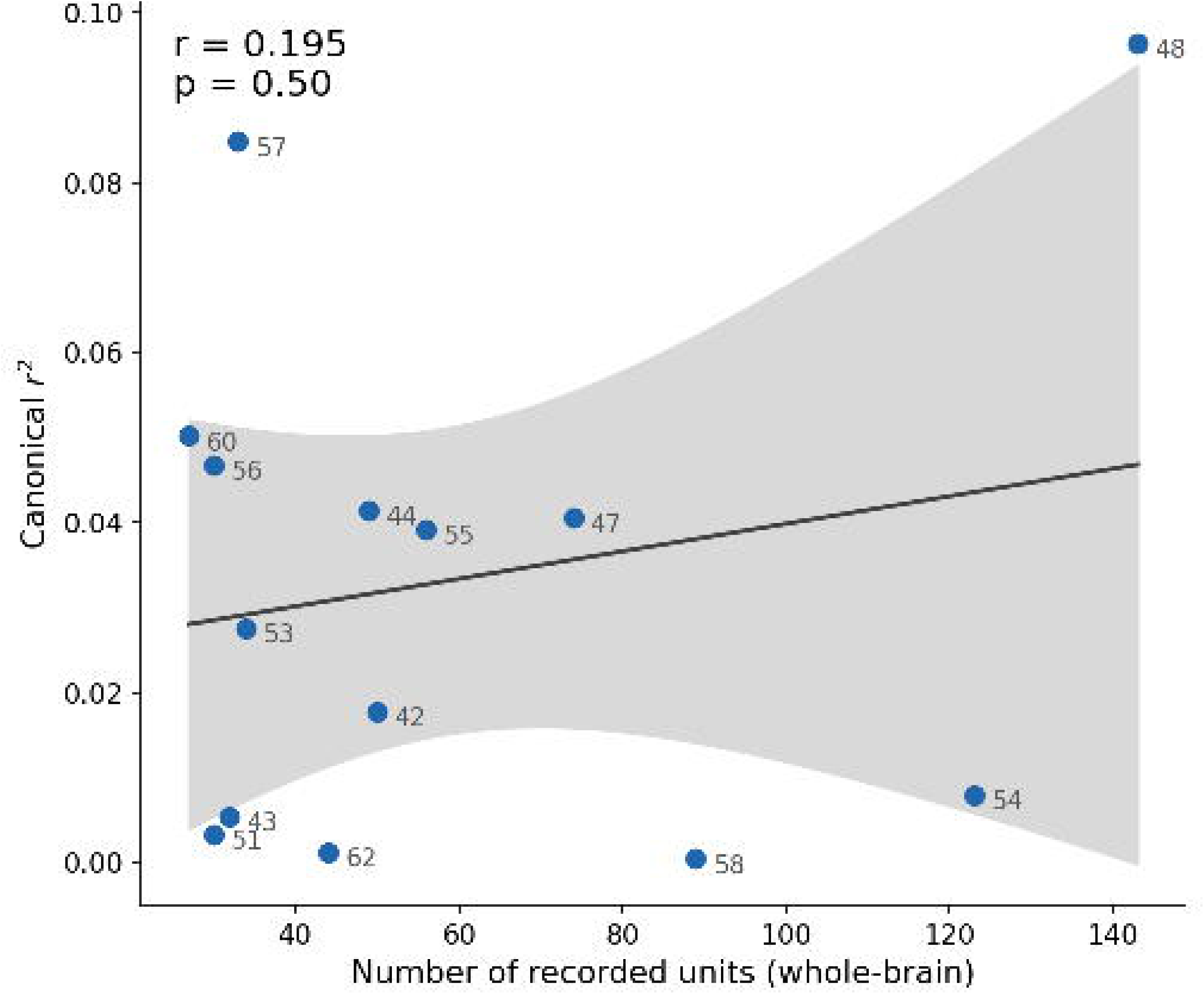
Population-size control. Per-subject canonical r² versus total recorded units (n = 14); solid line = linear fit, shaded band = 95% confidence interval. Pearson r = 0.196, p = 0.50 (two-tailed). Coupling strength shows no significant scaling with the number of recorded neurons.

## 4. Discussion

We asked whether the distributed single-neuron population and the joint eye-movement state are coupled during naturalistic movie viewing, and how any such coupling is organized. We found a group-level coupling (mean held-out canonical r = 0.145) that was individually significant in 9 of 14 subjects. The coupling was carried principally by saccade dynamics and pupil size, with fixation contributing only marginally. Critically, three convergent analyses (i.e. region loadings, leave-one-region-out, and single-neuron univariate tests) showed the coupling to be distributed and redundant: it is spread across medial-frontal and limbic populations, no region is strictly necessary, and almost no individual neuron survives correction.

This organization reframes what it means for the brain to track the eye-movement state. After correction, individual cells barely exceeded chance. Yet the same weakly coupled cells, considered jointly, cohere into a single, reliable population dimension aligned with the eye-movement state. The coupling to the eye-movement state is thus a low-dimensional, population-level property.

The anatomy of the coupling is interpretable. Saccades, the dominant carrier feature, leaned on medial-frontal cortex (ACC and preSMA) in the leave-one-region-out analysis, regions long implicated in the control and monitoring of eye movements and in salience-driven orienting, although at the single-neuron level saccade coupling was elevated across all sampled regions rather than focal (Corbetta and Shulman, 2002; Krauzlis et al., 2013). Pupil size, the second feature, showed its strongest single-neuron coupling in ACC, consistent with the tight link between pupil-linked arousal, ascending neuromodulation and cingulate activity (Aston-Jones and Cohen, 2005; Joshi et al., 2016; McGinley et al., 2015). Fixation duration, the weakest carrier, drew on the limbic amygdala and hippocampus, in accordance with a role for medial-temporal structures in memory-guided and value-guided viewing during naturalistic exploration. These regional emphases raise a question the studies above cannot settle: is each region specialized for a different aspect of eye behavior, or is there a single coordinated state expressed unevenly across the brain? A design that samples one region and one eye variable at a time cannot distinguish the two. Recording five regions simultaneously and treating the eye variables as a joint state can, and our data favor the second account on three counts. The three eye variables load onto one population dimension rather than onto separate regional ones. No single region’s removal significantly reduced that dimension. And single-neuron saccade coupling was elevated in every region we sampled rather than concentrated in medial-frontal cortex. The regional weights are therefore better read as gradations within one distributed dimension than as evidence for a dedicated oculomotor locus.

Functional MRI has repeatedly reported distributed rather than focal correlates of eye behavior: pupil fluctuations covary with antagonistic large-scale cortical systems (Yellin et al., 2015) and with the salience network, thalamus and frontoparietal cortex (Schneider et al., 2016); blinks shift activity between default-mode and dorsal-attention networks (Nakano et al., 2013); fixation duration loads on visual, prefrontal and medial superior frontal areas (Henderson and Choi, 2015); and saccades engage a frontal–parietal network (Grosbras et al., 2005). Our result is consistent with that picture and resolves an ambiguity in it. Because a voxel pools hundreds of thousands of neurons, a distributed BOLD correlate is equally compatible with a few strongly coupled cells diluted by averaging and with many weakly coupled ones. Here almost no neuron survives correction, yet the same cells read out jointly yield a reliable dimension: the distribution is genuine. The measurement also differs in kind: the coupling retains fluctuations up to roughly 0.5 Hz, several times faster than the hemodynamic response, and spike counts avoid the vascular contribution that pupil-linked arousal makes to BOLD (Chang et al., 2016). Our findings thus extend to the human brain, and to deep medial-frontal and limbic structures, the account from rodent brain-wide recordings in which spontaneous behavior explains much of population activity (Stringer et al., 2019; Musall et al., 2019).

Beyond the specific result, the study offers a methodological template. Coupling between slow physiological signals is easily explained by shared drift and by analytic leakage across cross-validation boundaries; we guarded against both with per-block linear detrending, contiguous block cross-validation, and a circular-shift null that preserves the autocorrelation and cross-feature covariance of the behavioral signals. The reliability of the effect across subjects, its insensitivity to neuron count, and its survival of these controls together argue that it reflects genuine neural–behavioral alignment rather than a drift or sampling artifact.

However, even though the results stand the above-mentioned control analyses, there are several limitations to keep in mind. First, the analysis is correlational: it establishes a shared dimension but not its direction or causal origin. Second, the recordings come from patients with epilepsy and from electrodes placed on clinical grounds; the sampled population spans five medial-frontal and limbic regions, so “distributed” here means distributed across the recorded regions rather than across the whole brain; visual, parietal and frontal oculomotor areas were not sampled, so the reported regional emphasis is conditioned on the available coverage. Additional work should test causal and directional structure, extend coverage to visual and oculomotor cortex, relate the coupling to moment-to-moment movie content and salience, and seek replication across stimuli and datasets.

In summary, the moment-to-moment eye-movement state is reflected in the coordinated activity of human single-neuron populations during naturalistic viewing. The signal is carried mainly by saccades and pupil size, it is distributed redundantly across medial-frontal and limbic populations and is recoverable only when the population is read out jointly. Eye movements thus relate to a coordinated, population-level neural state: a compact demonstration of how eye-tracking can serve as a variable for exploring human neural circuitry.

## CRediT author contributions

Carlo Cerquetella: Conceptualization, Methodology, Software, Formal analysis, Investigation, Data curation, Writing – original draft, Visualization. Salman E. Qasim: Conceptualization, Methodology, Formal analysis, Supervision, Funding acquisition, Writing – review and editing.

## Ethics

This study is a secondary analysis of the publicly available, de-identified dataset DANDI:000623 (Keles et al., 2024). The original recordings were obtained under the ethical approvals and written informed consent reported in that publication; no new data were collected for the present work.

## Data and code availability

The dataset analyzed here is publicly available as DANDI:000623 (Keles et al., 2024; CC-BY-4.0). Analysis code supporting the findings is available at https://github.com/CarloCerquetella/Neuropsychologia_SpecialIssue_EyeBrain (repository will be public upon publication).

## Supplementary material

**Supplementary Fig. S1.** (A) Neural and eye canonical variates across the movie; thick lines show the slow component (>30 s), which is strongly shared (r = 0.84). Interleaved cross-validation interpolates across this trend and counts it as coupling; dashed lines mark the five contiguous cross-validation blocks, gold shading the block zoomed below. (B) Within a single held-out block (∼96 s), a residual shared slow component remains (r = 0.74) that block cross-validation alone does not remove; this is what the detrend targets. (C) The same block after detrending: the fast (<30 s) canonical variates, projected from a fit on the other four blocks, still track one another (block r = 0.33). Panels use the global Gaussian high-pass variant (σ = 30 s) so that slow and fast components can be separated explicitly, for which this subject’s held-out r = 0.281; the headline analysis instead applies a per-block linear detrend, giving r = 0.311 for this subject (Table 1). Canonical variates are from a full, non-cross-validated fit and are illustrative only, all inferential statistics come from the held-out, circular-shift-tested analysis.

**Table S1.** Per-subject weights of the eye-side canonical variate (full fit, global Gaussian high-pass; normalized to unit total absolute weight). Held-out canonical r and significance from the headline analysis are repeated for reference.

| Subject | r | Sign | Pupil | Saccade | Fixation |
| --- | --- | --- | --- | --- | --- |
| CS42 | 0.133 | * | -0.773 | 0.220 | 0.007 |
| CS43 | -0.073 |  | 0.106 | 0.628 | 0.267 |
| CS44 | 0.204 | * | -0.042 | 0.682 | 0.276 |
| CS47 | 0.201 | * | -0.083 | 0.831 | -0.086 |
| CS48 | 0.311 | * | -0.008 | 0.942 | 0.050 |
| CS51 | 0.057 |  | -0.196 | 0.689 | -0.115 |
| CS53 | 0.166 | * | -0.723 | 0.108 | -0.170 |
| CS54 | 0.089 |  | -0.647 | 0.329 | 0.024 |
| CS55 | 0.198 | * | -0.616 | 0.271 | 0.113 |
| CS56 | 0.216 | * | -0.171 | 0.598 | 0.230 |
| CS57 | 0.291 | * | -0.192 | 0.750 | -0.058 |
| CS58 | -0.020 |  | -0.656 | 0.090 | 0.254 |
| CS60 | 0.224 | * | -0.227 | 0.638 | 0.135 |
| CS62 | 0.035 |  | 0.517 | 0.238 | -0.245 |

**Table S2.** Per-subject, per-feature attribution (held-out single-feature canonical correlation; empirical p against the circular-shift null).

| Subject | Pupil r | Pupil p | Sig Pupil | Saccade r | Saccade p | Sig Saccade | Fix r | Fix p | Sig Fix |
| --- | --- | --- | --- | --- | --- | --- | --- | --- | --- |
| CS42 | 0.149 | 0.006 | * | 0.112 | 0.034 | * | 0.050 | 0.186 |  |
| CS43 | 0.084 | 0.112 |  | 0.011 | 0.519 |  | -0.033 | 0.675 |  |
| CS44 | -0.002 | 0.459 |  | 0.225 | 0.002 | ** | 0.080 | 0.060 |  |
| CS47 | 0.068 | 0.118 |  | 0.216 | 0.002 | ** | 0.082 | 0.080 |  |
| CS48 | 0.103 | 0.038 | * | 0.337 | 0.002 | ** | -0.032 | 0.719 |  |
| CS51 | -0.009 | 0.435 |  | 0.087 | 0.088 |  | 0.098 | 0.020 | * |
| CS53 | 0.183 | 0.002 | ** | 0.011 | 0.493 |  | 0.007 | 0.461 |  |
| CS54 | 0.116 | 0.024 | * | 0.086 | 0.072 |  | 0.024 | 0.321 |  |
| CS55 | 0.226 | 0.004 | ** | 0.151 | 0.004 | ** | 0.187 | 0.002 | ** |
| CS56 | 0.211 | 0.002 | ** | 0.162 | 0.002 | ** | -0.008 | 0.511 |  |
| CS57 | 0.133 | 0.032 | * | 0.286 | 0.002 | ** | 0.096 | 0.034 | * |
| CS58 | 0.026 | 0.329 |  | 0.043 | 0.232 |  | 0.006 | 0.475 |  |
| CS60 | -0.034 | 0.699 |  | 0.232 | 0.002 | ** | 0.052 | 0.166 |  |
| CS62 | 0.058 | 0.208 |  | 0.019 | 0.367 |  | -0.033 | 0.713 |  |

**Table S3.** Leave-one-region-out per feature (uncorrected). Saccade leans on ACC and preSMA; fixation leans on amygdala and hippocampus.

| Region | Feature | N | r (full) | r (drop) | $\Delta r$ | t p (raw) |
| --- | --- | --- | --- | --- | --- | --- |
| ACC | pupil | 12 | 0.097 | 0.083 | 0.013 | 0.159 |
| ACC | saccade | 12 | 0.146 | 0.115 | 0.031 | 0.033 * |
| ACC | fixation | 12 | 0.044 | 0.044 | -0.001 | 0.522 |
| amygdala | pupil | 12 | 0.087 | 0.082 | 0.005 | 0.342 |
| amygdala | saccade | 12 | 0.150 | 0.142 | 0.007 | 0.263 |
| amygdala | fixation | 12 | 0.051 | 0.033 | 0.018 | 0.041 * |
| hippocampus | pupil | 12 | 0.098 | 0.098 | 0.000 | 0.490 |
| hippocampus | saccade | 12 | 0.163 | 0.160 | 0.002 | 0.349 |
| hippocampus | fixation | 12 | 0.053 | 0.036 | 0.018 | 0.114 |
| preSMA | pupil | 13 | 0.095 | 0.079 | 0.016 | 0.091 |
| preSMA | saccade | 13 | 0.151 | 0.120 | 0.031 | 0.043 * |
| preSMA | fixation | 13 | 0.047 | 0.041 | 0.006 | 0.335 |
| vmPFC | pupil | 10 | 0.115 | 0.115 | 0.000 | 0.497 |
| vmPFC | saccade | 10 | 0.151 | 0.158 | -0.007 | 0.915 |
| vmPFC | fixation | 10 | 0.034 | 0.042 | -0.008 | 0.914 |

**Table S4.**
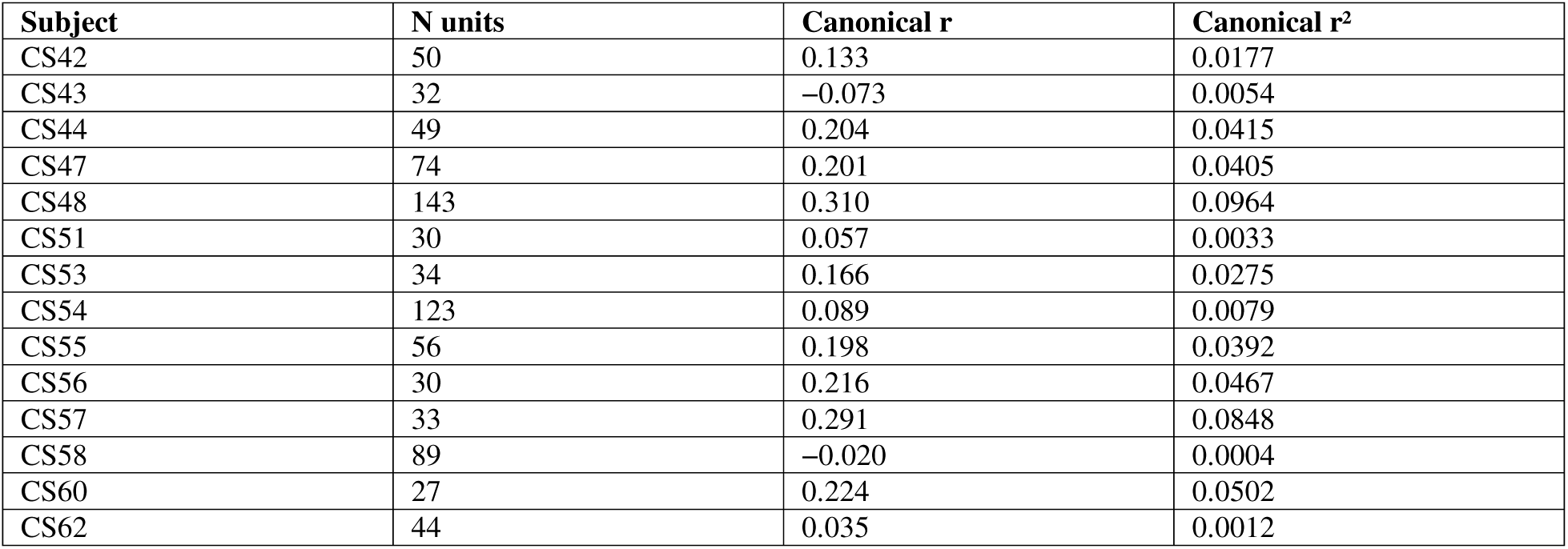
Per-subject canonical r and r² (used in the population-size control, Fig. 7).

