## Supplementary figures and images for "Human single-neuron recordings reveal population coding of attentional dynamics during naturalistic movie viewing"

### Supplementary Figure 1

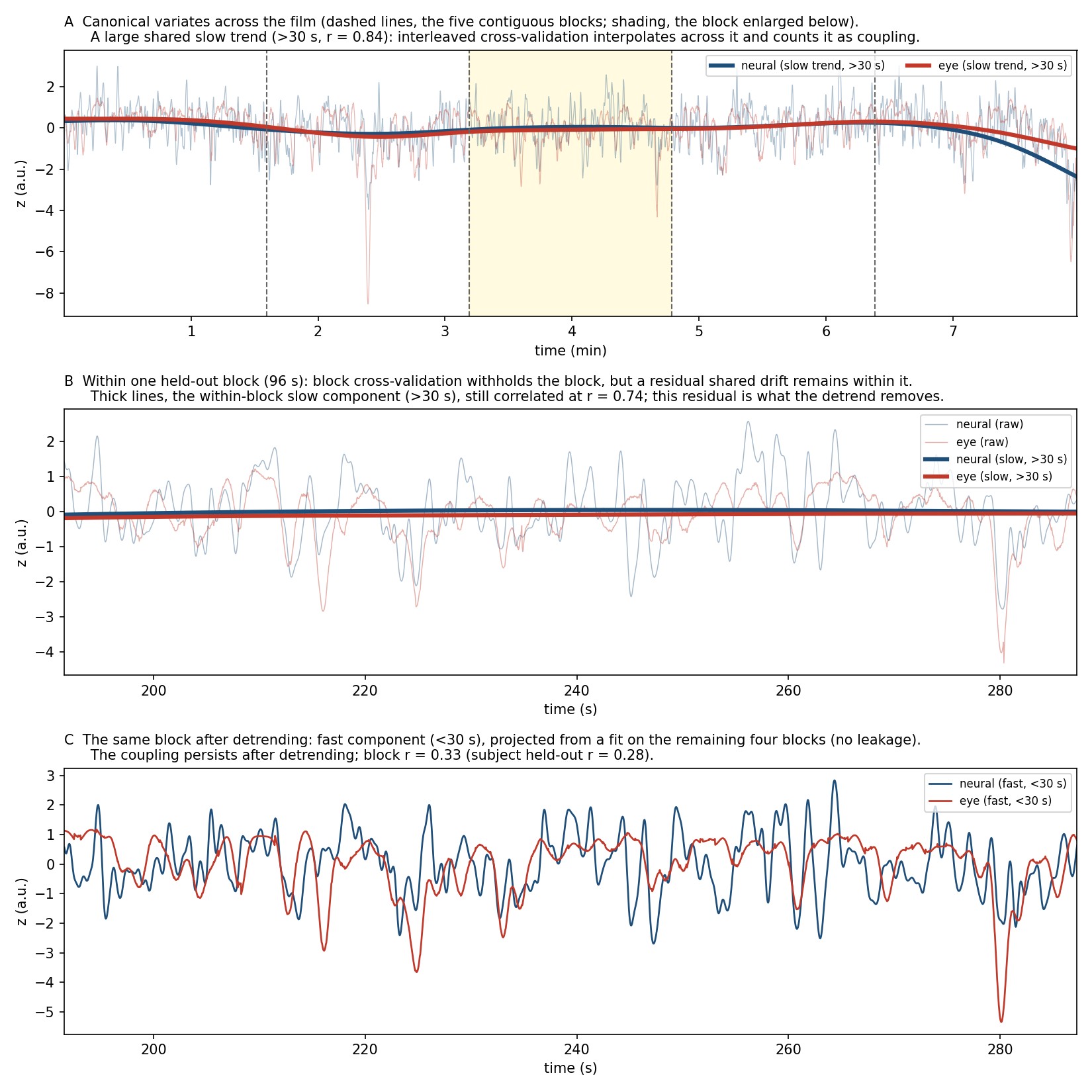
